# Advanced age attenuates the antihyperalgesic effect of morphine and decreases μ-opioid receptor expression and binding in the rat midbrain Periaqueductal Gray in male and female rats

**DOI:** 10.1101/2020.06.16.154740

**Authors:** Evan F. Fullerton, Myurajan Rubaharan, Mary C. Karom, Richard I. Hanberry, Anne Z. Murphy

## Abstract

The present study investigated the impact of advanced age on morphine modulation of persistent inflammatory pain in male and female rats. The impact of age, sex, and pain on μ-opioid receptor (MOR) expression and binding in the ventrolateral PAG (vlPAG) was also examined using immunohistochemistry and receptor autoradiography. Intraplantar administration of Complete Freund’s adjuvant induced comparable levels of edema and hyperalgesia in adult (2-3mos) and aged (16-18mos) male and female rats. Morphine potency was highest in adult males, with a two-fold decrease in morphine EC_50_ observed in aged versus adult males (10.22mg/kg versus 5.19mg/kg). Adult and aged female rats also exhibited significantly higher EC_50_ values (10.69 mg/kg and 9.00 mg/kg, respectively) compared to adult males. The upward shift in EC_50_ from adult to aged males was paralleled by a reduction in vlPAG MOR expression and binding. The observed age-related reductions in morphine potency and vlPAG MOR expression and binding have significant implications in pain management in the aged population.

## 1.1 Introduction

The United States population is rapidly aging, and in the coming decade, over 20% of US citizens will be 65 years of age or older (CDC, 2013). It is estimated that over 50% of these individuals will experience persistent and/or severe pain, including pain associated with arthritis, cancer, diabetes mellitus, and cardiovascular disease (Robeck et al., 2014). Persistent pain in the elderly typically involves multiple sites and is comorbid with a variety of other conditions, resulting in depression, sleep disruption, cognitive impairment, and decreased socialization. Together, these factors potentially decrease the quality of life and increase the risk of early death (Makris et al., 2015; Reid et al., 2015; Patel et al., 2013; Domenichiello and Ramsden, 2019). Although opioids, including morphine and fentanyl, are the most potent and commonly prescribed analgesics for chronic pain management, pain in the elderly is typically undermanaged or ignored altogether (Pergolizzi et al. 2008). Several factors contribute to the undermanagement of pain in the elderly, including under-reporting of pain intensity by the patient, physician reluctance to prescribe opioids due to fears of adverse side effects including cognitive impairment and respiratory depression, and misconceptions on the part of the patient and physician regarding tolerance and addiction (Cavalieri, 2005; Naples et al., 2016; Prostran et al., 2016; Saunders et al. 2010; Miaskowski, 2011; Miaskowski et al., 2011 Schiltenwolf et al., 2014).

Although opioids for chronic pain management in the elderly are finally beginning to increase (Pokela et al., 2010; Lapane et al., 2013), there remains a dearth of information regarding the suitable dosing regimen (Campbell et al., 2010). Clinical studies examining opioid potency in the elderly are inconsistent, with reports of increased, decreased, or equivalent sensitivity compared to adults (Kaiko, 1980; Gagliese and Katz, 2003; Papaleontiou et al. 2010; Prostran et al., 2016). Several factors likely contribute to these different results. First, although opioids are prescribed for the alleviation of severe and persistent pain, the majority of these studies were conducted in elderly patients experiencing acute pain. The presence of comorbid conditions such as diabetes and high blood pressure, and the use of concomitant medications, may also either augment or potentiate morphine’s analgesic effects. These studies also typically fail to include sex as a biological variable, either not reporting it or focusing exclusively on males (Gagliese and Katz, 2003; Aubrun et al., 2005; Lautenbacher, 2012), leaving the impact of age on opioid modulation of pain in women largely unknown. Sex and age differences in the likelihood of self-reporting pain in a clinical setting, along with the subjective nature of pain, further contribute to the contradictory findings regarding effective dosing in the aged population (Ferrell et al., 1991; Reddy et al., 2012; Dampier et al., 2013).

Preclinical studies in rodents have also failed to provide insight regarding the impact of advanced age on opioid modulation of pain. First, very few studies have been conducted, and similar to their clinical counterparts, these studies report increased, decreased, and no change in opioid potency as a function of age (Chan and Lai, 1982; Kramer and Bodnar, 1986; Gagliese and Melzack, 2000). These studies also suffer from the same shortfalls identified for clinical studies, including acute rather than chronic pain assays (typically tail-flick or hot plate) and use of male subjects exclusively (Chan and Lai, 1982; Kavaliers et al., 1983; Smith and Gray, 2001; Jourdan et al., 2002). Therefore, a clear picture regarding the impact of advanced age on opioid potency has yet to emerge.

The endogenous descending analgesia circuit, consisting of the midbrain periaqueductal gray (PAG) and its descending projections to the rostral ventromedial medulla (RVM) and dorsal horn of the spinal cord, is also a critical neural circuit for exogenous pain modulation (Bausbaum et al., 1978; Bausbaum and Fields, 1979; Bausbaum et al., 1976; Morgan et al., 1992, Morgan et al., 2006). The ventrolateral PAG (vlPAG) contains a large population of mu-opioid receptor (MOR) containing neurons, the preferred receptor for morphine (Martin, 1963; Wolozin and Pasternak, 1981), and direct administration of MOR agonists into the PAG produces potent analgesia (Satoh et al., 1983; Jensen et al., 1986; Bodnar et al., 1988). Conversely, direct administration of MOR antagonists into the PAG, or lesions of PAG MOR, significantly attenuate the analgesic effect of systemic morphine (Wilcox et al., 1979; Ma and Han, 1991; Zhang et al., 1998; Loyd et al., 2008). Despite high rates of chronic pain among the elderly, and reported age differences in morphine potency, surprisingly little is known regarding how advanced age influences the distribution and function of MOR in the PAG.

Here we present a series of experiments delineating the impact of advanced age on opioid analgesia and the expression and binding of MOR in the vlPAG of the male and female rat. These studies provide the first data on age-mediated changes in the cellular and molecular processes of a central neural circuit governing pain and opioid analgesia using a clinically relevant pain assay.

## 2 Materials and methods

### 2.1 Experimental subjects

Adult (2-3mos) and aged (16-18mos) male and regularly cycling female Sprague–Dawley rats were used in these experiments (Charles River Laboratories, Boston, MA). Rats were co-housed in same-sex pairs on a 12:12 hour light/dark cycle (lights on at 08:00 am). Access to food and water was ad libitum throughout the experiment, except during testing. All studies were approved by the Institutional Animal Care and Use Committee at Georgia State University and performed in compliance with Ethical Issues of the International Association for the Study of Pain and National Institutes of Health. All efforts were made to reduce the number of rats used in these experiments and to minimize pain and suffering.

### 2.2 Vaginal cytology

Beginning ten days prior to testing, vaginal lavages were performed daily on adult and aged female rats to confirm that all rats were cycling regularly and to keep daily records of the stage of estrous. Proestrus was identified as a predominance of nucleated epithelial cells, and estrus was identified as a predominance of cornified epithelial cells. Diestrus 1 was differentiated from diestrus 2 by the presence of leukocytes. Rats that appeared between phases were noted as being in the more advanced stage (Loyd et al., 2007).

### 2.3 Behavioral testing

Adult (2-3mos) and aged (16-18mos), male and female rats were used in the behavioral studies examining the impact of age and sex on morphine potency (n=6-10/group; N=32). As previously described, thermal nociception was assessed using the paw thermal stimulator (Univ. California San Diego) (Hargreaves et al., 1988; Wang et al., 2006). Briefly, the rat was placed in a clear plexiglass box resting on an elevated glass plate maintained at 30°C. A timer was initiated as a radiant beam of light was positioned under the hind paw, and the time point at which the rat retracted its paw in response to the thermal stimulus was electronically recorded as the paw withdrawal latency (PWL) in seconds (s). A maximal PWL of 20.48s was used to prevent tissue damage due to the repeated application of a noxious thermal stimulus. Rats were acclimated to the testing apparatus 30–60 min before the start of the experiment on the day of testing. All behavioral testing took place between 08:00 am and 12:00 pm (lights on at 08:00 am). The maximum temperature of the thermal stimulus was recorded before and after each trial to maintain consistent recordings between groups and did not exceed a range of 60–65°C throughout the experiments. All testing was conducted blind with respect to group assignment.

### 2.4 Inflammatory hyperalgesia and edema

Following baseline PWL determination, persistent inflammation was induced by injection of complete Freund's adjuvant (CFA; Mycobacterium tuberculosis; Sigma; 200μl), suspended in an oil/saline (1:1) emulsion, into the plantar surface of the right hind paw. Paw diameters were determined using calibrated calipers applied midpoint across the plantar surface of both hind paws before and after induction of inflammation.

### 2.5 Morphine administration

Twenty-four hours following CFA administration, rats were administered morphine using a cumulative dosing paradigm as previously described (Loyd et al. 2008). Briefly, rats received subcutaneous injections of morphine sulfate every 20 minutes (1.8, 1.4, 2.4, 2.4, 2.0), resulting in the following doses: 1.8, 3.2, 5.6, 8.0, and 10 mg/kg (s.c; NIDA; Bethesda, MD, USA). PWLs were determined 15 minutes after each administration. Morphine sulfate was prepared in a saline vehicle within 24 hours of administration.

### 2.6 Perfusion fixation

Immunohistochemical localization of MOR was determined in a separate cohort of adult and aged male and female rats. To determine the impact of persistent inflammatory pain on MOR expression, a subset of rats in each treatment group received an intraplantar injection of CFA 24-72 hrs prior to perfusion (total of 8 treatment groups; n=5-8; N =45). Rats were given a lethal dose of Euthanasol (i.p.) and transcardially perfused with 200-250 mL of 0.9% sodium chloride containing 2% sodium nitrite as a vasodilator to remove blood from the brain, followed by 300-400 mL of 4% paraformaldehyde in 0.1 M phosphate as a fixative. Immediately following perfusion, brains were removed and stored in 4% paraformaldehyde solution. After 24 hours, brains were placed in a 30% sucrose solution and stored at 4°C for at least one week prior to sectioning. Brains were cut coronally into 25-μm sections with a Leica 2000R freezing microtome and stored free-floating in cryoprotectant-antifreeze solution (Watson et al., 1986) at −20 °C until immunohistochemical processing.

### 2.7 Immunohistochemistry

Coronal sections were removed from cryoprotectant and processed for MOR immunoreactivity, as previously described (Loyd et al., 2008). Briefly, sections were rinsed extensively in potassium phosphate-buffered saline (KPBS) to remove the cryoprotectant solution. After rinsing in KPBS, sections were incubated in a blocking solution of 5% Normal Donkey Serum in KPBS containing 0.5% Triton X-100 for 1 h at room temperature. The tissue was then incubated in an antibody solution of rabbit anti-MOR (specifically targeting the extracellular N-terminus of the receptor; Alomone Labs, Jerusalem, Israel; 1:1,000) in KPBS containing 1.0% Triton X-100 for 1 h at room temperature followed by 48h at 4°C. After rinsing in KPBS, sections were incubated in a Donkey anti-Rabbit 488 (1:200) solution for 2h at room temp. Tissues were then rinsed using KPBS and mounted using SlowFade Diamond Antifade Mountant (Invitrogen). Images were processed on Zeiss LSM 700 Confocal Microscope at 40x.

To quantify neuronal number within the vlPAG, a second set of sections were stained using mouse anti-NeuN (Millipore Bioscience Research Reagents; 1:50,000). Following incubation in primary antibody solution as described above, the tissue was rinsed with KPBS, incubated for 1h in biotinylated goat anti-mouse IgG (Jackson Immunoresearch, West Grove, PA, USA; 1:200), rinsed again with KPBS, and then incubated for 1 h in avidin-biotin-peroxidase complex (1:10; ABC Elite Kit, Vector Laboratories). After rinsing in KPBS and sodium acetate (0.175 M; pH 6.5), NeuN immunoreactivity was visualized as a black reaction product using nickel sulfate intensified 3,31-diaminobenzidine containing 0.08% hydrogen peroxide in KPBS (pH 7.4). After incubating for 15–30 min, the reaction was terminated by rinsing in sodium acetate buffer. Tissue sections were mounted out of KPBS onto gelatin-subbed slides, air-dried overnight, dehydrated in a series of graded alcohols, cleared in xylene, and coverslipped using Permount.

### 2.8 Anatomical data analysis and presentation

Fluorescent images were captured on Zeiss LSM 700 Confocal Microscope at 40x, and MOR intensity (signal intensity/signal volume) was calculated using Imaris software. MOR intensity values were determined for the left and right ventrolateral subdivision of each PAG image from two representative levels of the mid-caudal PAG (Bregma −7.74 and −8.00) and were averaged for each slice. Data were analyzed across two representative levels of the mid-caudal PAG (Bregma −7.74 and −8.00).

Photomicrographs from Ni-DAB stained sections were generated using a Synsys digital camera attached to a Nikon Eclipse E800 microscope. Images were captured with IP Spectrum software and densitometry of labeling assessed in ImageJ. Densitometry measurements were conducted bilaterally as separate images of the left and right side and averaged. As there was no significant effect of rostrocaudal level in the analyses, data are collapsed and presented as vlPAG. MOR intensity and NeuN densitometry values are expressed as the mean ± standard error of the mean (SEM). Previous data have shown that there are no sex differences in total area (mm^2^) of the PAG between male and female Sprague–Dawley rats (Loyd & Murphy, 2006). All images were collected and analyzed by an experimenter blinded to the experimental condition.

### 2.9 Receptor autoradiography

To determine the impact of age, sex, and pain on vlPAG MOR binding, a separate cohort of adult and aged, male and female, handled and CFA treated rats (total of 8 treatment groups; n=6; N=48) were used for receptor autoradiography. Rats were restrained using DecapiCones and decapitated. Brains were removed rapidly, flash-frozen in 2-methyl butane on dry ice, and stored at −80°C. Frozen tissue was sectioned in a 1:6 series of 20μm coronal sections at −20°C with a Leica CM3050S cryostat. Sections were immediately mounted onto Super-frost slides (20°C) and stored at −80°C until the time of the assay. Tissue was processed for autoradiography as previously described (Loyd et al., 2008; LaPrairie and Murphy, 2009). Briefly, sections were allowed to thaw to room temperature and fixed in 0.1% paraformaldehyde followed by rinses in 50mM Tris buffer, pH 7.4. Slides were then placed in a tracer buffer containing tritiated DAMGO (100 nm; American Radiolabeled Chemicals) for 60 min followed by a series of rinses in 50 mm Tris buffer, pH 7.4, containing MgCl_2_. Following a final dip in cold dH_2_0, tissue was allowed to dry at room temp. Slides were then apposed to FujiFilm imaging plates along with [^3^H]-microscale standards (Perkin-Elmer/NEN, MA, USA) for six weeks. Image plates were processed with a FujiFilm BAS 5000. Autoradiographic [^3^H]-receptor binding was quantified from the images using Scion Image software. [^3^H]-standards were used to convert uncalibrated optical density to disintegrations per minute (DPM). For analysis, DPM values were determined for the left and right ventrolateral subdivision of each PAG image from two representative levels of the mid-caudal PAG (Bregma −7.74 and −8.00) and averaged for each slice.

### 2.10 Statistical Analysis and Data Presentation

All values are reported as mean ± SEM. For behavioral data analysis, data are expressed as either raw PWLs or percent maximal possible effect (%MPE), defined as [(PWL – CFA baseline)/(maximal PWL – CFA baseline)] × 100. Significant main effects of sex, age, and treatment were assessed using ANOVA or Repeated Measures ANOVA; p < 0.05 was considered statistically significant. Tukey’s post-hoc tests were conducted to determine significant mean differences between groups that were apriori specified. Our behavioral data suggested that morphine was less potent in aged males and females, leading us to hypothesize that MOR expression and binding were reduced in the vlPAG of aged rats as well as females; thus, one-tailed post-hoc tests were used for immunohistochemistry and autoradiography. Anatomical data are expressed either as percent area covered for Ni-DAB IHC, intensity (signal intensity/signal volume) for fluorescent IHC, or disintegrations per minute (DPM) for autoradiography data. For data presentation, a representative animal from each experimental group was selected, and photomicrographs generated using a confocal microscope. Images were captured and processed with Imaris software (immunohistochemistry) or processed with Scion Image (autoradiography). Alterations to the images were strictly limited to the enhancement of brightness and contrast.

## 3 Results

### 3.1 No impact of advanced age or sex on basal thermal nociception or CFA-induced hyperalgesia

To assess the impact of age and sex on baseline thermal nociception, paw withdrawal latencies (PWL) in response to a noxious thermal stimulus were determined (Hargreaves et al., 1988). No significant impact of age [F_(1,28)_ = 3.384, p = 0.078] or sex [F_(1,28)_ = 1.362, p = 0.253] was noted for baseline PWL (Figure 1A). Similarly, no significant effect of age [F_(1,33)_ = 2.323, p = 0.137] or sex [F_(1,33)_ = 2.533, p = 0.121] on CFA-induced hyperalgesia was observed (Figure 1B). In all 4 groups, CFA reduced PWL from 8.26s to 3.86s, an average % change in PWL of −53.4% (Figure 1C). The inflammatory insult induced by CFA injection resulted in comparable inflammation in all experimental groups, with no significant impact of age [F_(1,40)_ = 2.171, p = 0.149] or sex [F_(1,40)_ = 0.491, p = 0.487]. In all 4 groups, CFA increased paw diameter by an average % change of 168.2% (Figure 1D).

**Fig. 1.**
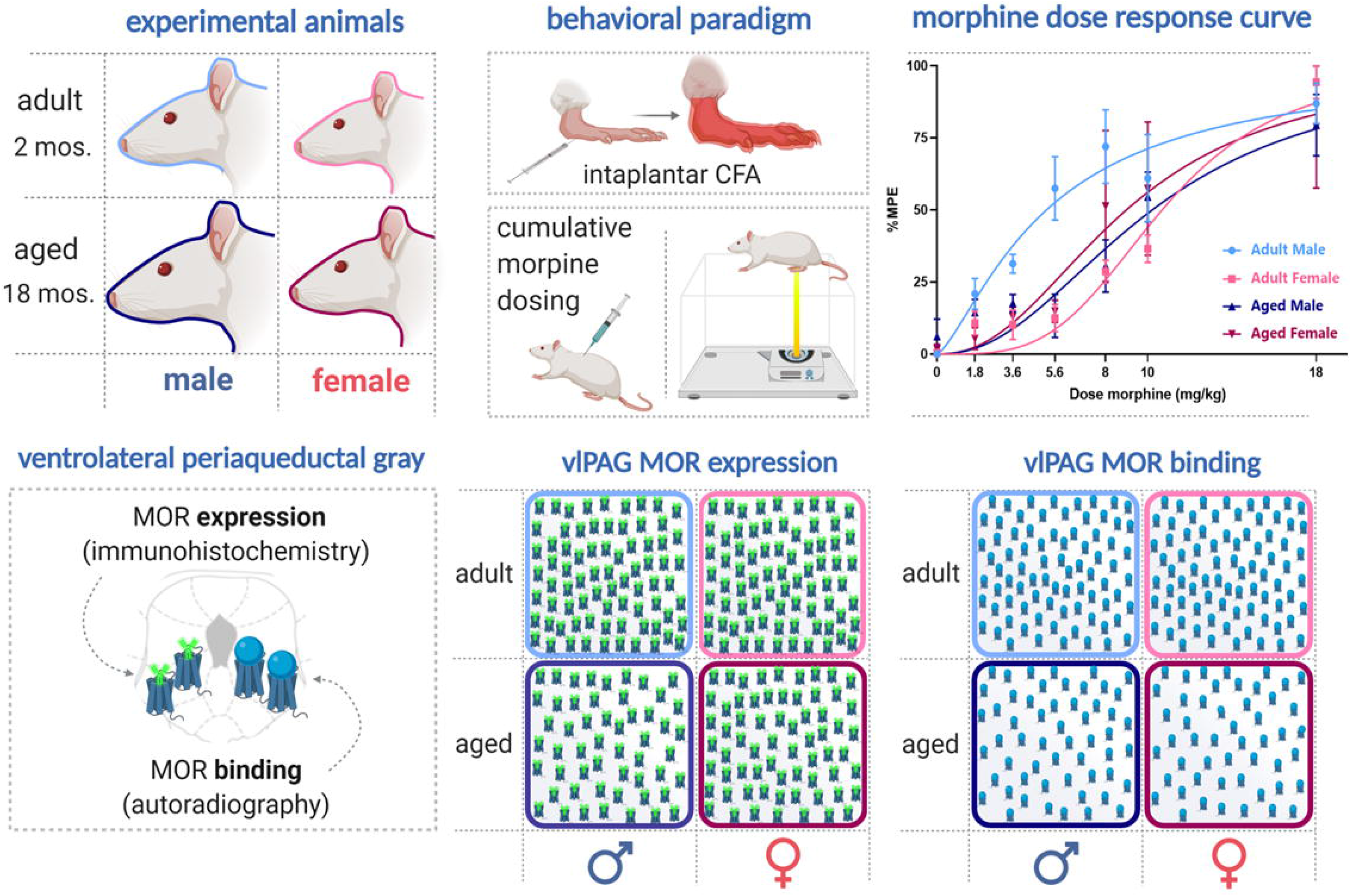
No impact of age or sex was observed for basal thermal nociception (**A**) or post-CFA PWLs (**B**). Similarly, no age or sex difference in CFA-induced hyperalgesia (**C**) or CFA-induced edema (**D**). Ns, not significant calculated by two-way ANOVA. Graphs indicate mean ± SEM.

### 3.2 Age and sex differences in morphine anti-hyperalgesia

We next determined the impact of age and sex on morphine potency using a cumulative dosing paradigm to derive EC_50_ (Figure 2A). Aged male rats were less sensitive to morphine than their adult counterparts, with a two-fold increase in EC_50_ compared to adult males (EC_50_ = 10.22 mg/kg vs. EC_50_=5.19). Adult females were also less sensitive to morphine than adult males (EC_50_ = 10.69 mg/kg vs EC_50_ = 5.19 mg/kg). Females showed no age differences, with similar EC_50_ values observed for adult and aged rats (EC_50_ 9.00mg/kg vs. EC_50_ 10.69) (Figure 2B). We also examined the impact of age and sex on %MPE at the 5.6 mg/kg dose (midpoint dose in the cumulative dosing paradigm). Results from a two-way ANOVA indicated a significant impact of age [F_(1,16)_ = 5.883, p = 0.028] and sex [F_(1,16)_ = 8.754, p = 0.009]. Post hoc analysis showed that morphine induced significantly greater anti-hyperalgesia in adult males than aged males (p = 0.0026), adult females (p = 0.0012), and aged females (p = 0.0076) (Figure 2C). Interestingly, morphine potency in aged males was similar to that for females, suggesting a similar attenuation in the mechanisms underlying opiate anti-hyperalgesia in females and aged males.

**Fig. 2.**
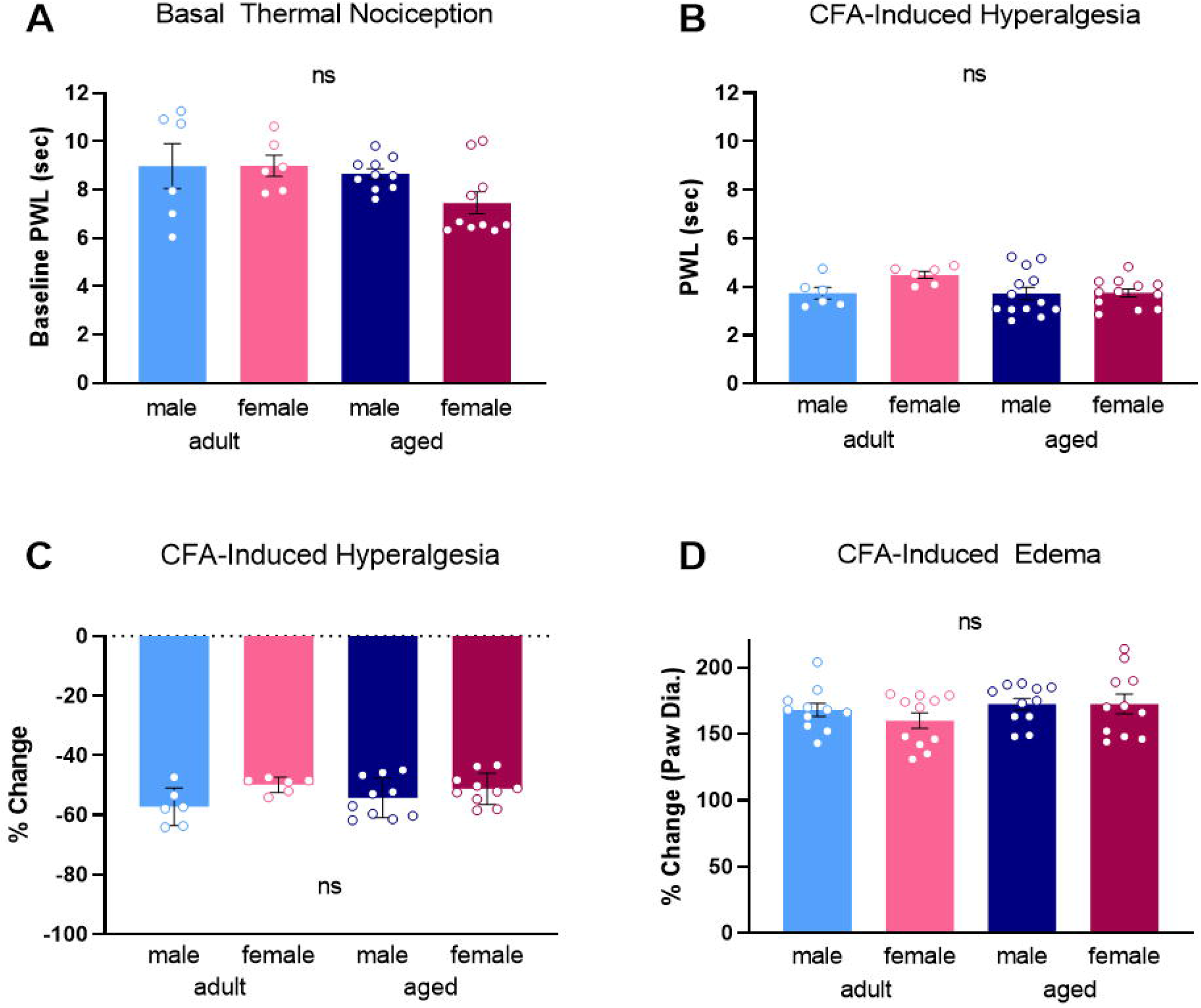
Morphine _EC_50__ values were generated from nonlinear regression analysis (**A**). Adult males exhibit lower morphine EC_50_ values than females and aged males (**B**). Adult males exhibit significantly higher morphine potency at 5.6mg/kg compared to females and aged males (**C**). Data presented in milligrams of morphine per kilogram of body weight. *Significant differences between adult male and adult female, adult male and aged male, and adult male and aged female. p<0.05 calculated by Tukey’s post hoc test. Graphs indicate mean ± SEM.

### 3.3 Impact of advanced age, sex, and chronic pain on MOR expression in the midbrain vlPAG

We next sought to determine if sex and age differences in mu-opioid receptor levels in the vlPAG contributed to our observed behavioral differences. An example of MOR immunostaining in the vlPAG is shown in Figure 3A. Our analysis indicated a significant impact of age on vlPAG MOR expression, with adult rats expressing higher MOR density [F_(1,37)_ = 9.933, p = 0.040] (Figure 3B). No significant impact of sex [F_(1,37)_ = 3.093, p = 0.243] or treatment [F_(1,37)_ = 0.855, p = 0.536] was observed. Post hoc analysis showed that vlPAG MOR expression was significantly greater in adult males compared to aged males (p = 0.013) (Figure 3B).

**Fig. 3.**
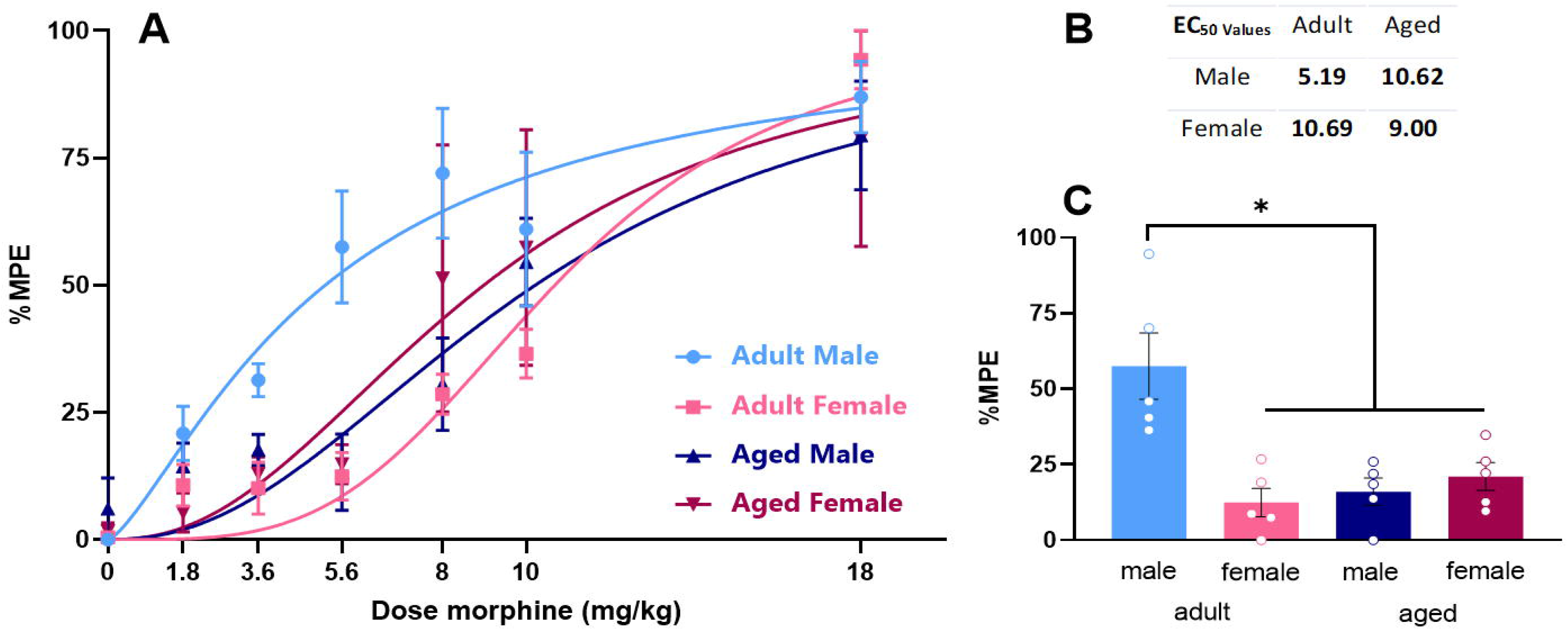
Representative section from rat brain with red highlight showing the vlPAG region quantified for MOR immunoreactivity (**A**). MOR densitometry in the vlPAG was significantly higher in adults compared to aged rats (**B**). No significant impact of sex or treatment were noted. No differences MOR density were observed between handled and CFA treated groups, so these data are combined. No significant effect of age or sex was observed in NeuN densitometry in the vlPAG (**C**). *Significant difference between adult and aged males; p<0.05 calculated by Tukey’s post hoc test. ns, not significant. Graphs indicate mean ± SEM.

To ensure that our observation of reduced vlPAG MOR in aged males was not due to an age-induced decrease in overall cell number, adjacent sections through the PAG were immunohistochemically stained using the neuron-specific marker NeuN (Millipore; [75]) and the density of staining quantified using densitometry. No significant effect of age [F_(1,23)_ = 0.133, p = 0.909] or sex [F_(1,23)_ = 0.072, p = 0.791] was observed in NeuN immunostaining (Figure 3C), indicating that the reduction in MOR density was not due to an overall age-related loss in neuronal number. Similarly, there was no significant effect of age [F_(1,10)_ = 2.146, p = 0.174] or sex [F_(1,10)_ = 0.083, p = 0.779] on MOR expression in the inferior colliculus (IC), a midbrain region rich in MOR, indicating the age-induced reduction in MOR expression was specific for the PAG (data not shown).

### 3.4 Impact of advanced age, sex, and chronic pain on MOR binding in the midbrain vlPAG

We next used autoradiography to determine if the observed reduction in MOR expression reflected a reduction in MOR binding. [^3^H]-DAMGO was used to label MOR as previously described (LaPrairie and Murphy, 2009) Representative autoradiograms are shown in Figure 4A. Consistent with what we noted using IHC, there was a significant impact of age [F_(1,39)_ = 30.08, p < .0001], with adults exhibiting overall greater binding than their aged counterparts (Figure 4B). Post hoc analysis showed that aged males had reduced MOR binding compared to adult males [p = 0.033]. There was no significant impact of sex [F_(1,39)_ = .108, p = 0.306] or treatment [F_(1,39)_ = 0.021, p = 0.885] (Figure 4B), and no significant interactions were noted.

**Fig. 4.**
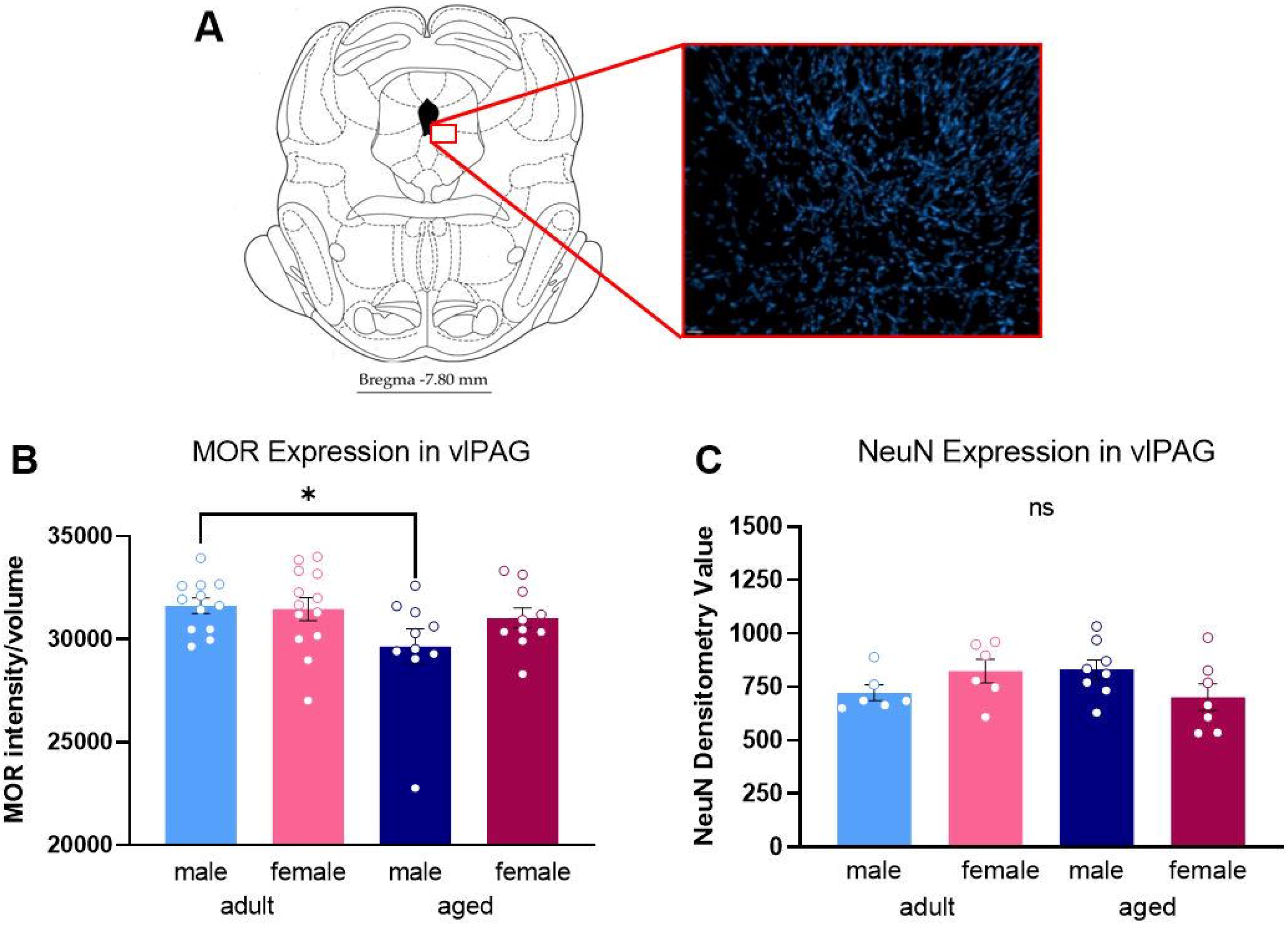
Heat mapped representative images of DAMGO binding in the midbrain of experimental groups (**A**). Aged rats exhibited reduced PAG MOR binding compared to adults (**B**). No impact of sex or treatment were found. No differences in density were observed between handled and CFA treated groups, so these data are combined. Data presented in disintegrations per minute per milligram. * Significant difference between adult males and females and aged rats; p<0.05 calculated by Tukey’s post hoc test. Graphs indicate mean ± SEM.

## Discussion

The present studies were conducted to determine the impact of advanced age on opioid modulation of pain. Using a clinically relevant model of persistent inflammatory pain, we report that aged rats require higher doses of morphine than their adult counterparts to achieve comparable levels of analgesia. The impact of age was sex specific, such that aged males required a significantly higher morphine dosage to reach analgesia compared to adult males; no impact of age was noted in morphine potency in females. Our results further demonstrate significant age differences in vlPAG MOR expression, with aged rats exhibiting reductions in vlPAG MOR protein and binding. As MOR expression in the vlPAG is critical for morphine attenuation of pain, the observed age-induced reduction in vlPAG MOR may drive the reduced morphine potency observed in aged male rats.

### No impact of advanced age or sex on baseline pain sensitivity and CFA-induced hyperalgesia

In the present studies, no effect of advanced age (or sex) was observed in baseline thermal sensitivity. This finding is consistent with our previous studies as well as those of others (Ali et al., 1995; Craft and Milholland, 1998; Craft et al., 1998; Wang et al., 2006). Indeed, the majority of rodent studies assessing the impact of sex on basal nociception report no differences (Wang et al., 2006; Mogil et al., 2000; Mogil et al., 2010; Racine et al., 2012; Mogil, 2020). Previous studies on age-associated changes in basal nociception are highly varied, with reports of increased (Hess et al., 1981; Kitagawa et al., 2005), decreased (Chan and Lai, 1982; Kramer et al., 1985; Garrison and Stucky, 2014; Muralidharan et al., 2020), and no change in mechanical or thermal pain sensitivity in aged rodents compared to adults (Crisp et al., 1994; Taguchi et al., 2010; Yezierski, 2012; Mecklenburg et al., 2017; Samir et al., 2017).

Similarly, meta-analyses of both clinical and laboratory measurements in humans fail to report a consistent impact of age on nociception (Gibson and Helme, 2001; Weyer et al., 2016; Ostrom et al., 2017). For example, in a meta-analysis summarizing the findings of 52 clinical research studies, Lautenbacher (2012) concluded that aged individuals exhibit increased pain thresholds (i.e., decreased sensitivity) compared to young adults. However, this data included both pain-free individuals and individuals who were experiencing chronic pain and did not include sex as a variable of analysis. A similar study by Ostrom et al., (2017) of over 3,400 participants reported no impact of age on pain sensitivity across numerous modalities (pressure, mechanical, thermal), although a limited age range was examined (18-44 years; Ostrom et al., 2017).

In the present study, intraplantar injection of the mycobacterium complete Freund's adjuvant was used to induce persistent inflammatory pain (Millan et al., 1988; Stein et al., 1988). We report no significant impact of either age or sex on the degree of hyperalgesia elicited by intraplantar CFA, or CFA-induced edema (Loyd et al., 2006; Wang et al., 2006; Loyd et al., 2008). To date, very few preclinical studies have examined the impact of age on persistent pain. Mecklenburg et al. (2017) reported no impact of age (3, 6, 18, and 24 months) on either mechanical or thermal hyperalgesia in male mice using a plantar incisional model of pain. Similar results were reported by Garrison and Stucky (2014) in male mice aged 3 and 24 months following intraplantar CFA. This study also reported comparable levels of edema in adult and aged mice up to 8 weeks post-CFA. In contrast, Weyer et al. (2016) reported that aged male mice (18 weeks) showed decreased mechanical hyperalgesia in comparison to adults across all eight weeks examined. This attenuated hyperalgesic response was accompanied by an overall reduction in noxious stimulus-evoked action potentials in the dorsal horn. Similar to the present study, no impact of age on CFA-induced edema was reported (Weyer et al., 2016).

### Impact of advanced age and sex on morphine analgesia

The present studies are the first to examine the impact of advanced age on opioid modulation of persistent inflammatory pain in male and female rats. We report morphine EC_50_ values dependent on age and sex; notably, aged male rats required significantly higher doses than their adult counterparts to produce comparable anti-hyperalgesia. Similar results were reported by Muralidharan et al. (2020), who noted a reduction in morphine's anti-allodynic effect in male mice 54 weeks of age. Interestingly, the anti-allodynic effects of the gabapentinoid pregabalin were also attenuated (Muralidharan et al., 2020), suggesting that the age-induced blunting of analgesics on pain are not limited to opioids, but rather, extend to other classes of drugs.

Clinically, several studies report that opioid requirements for post-operative pain are inversely related to age (Kaiko, 1980; Gagliese and Katz, 2003; Keita et al., 2009), although results to the contrary have also been reported (Papaleontiou et al. 2010; Prostran et al., 2016). Results from clinical studies on elderly individuals are more challenging to interpret due to the high degree of comorbidity with conditions such as heart disease, diabetes, and high blood pressure that may alter the perception of pain and necessitate the use of concomitant medications. Additional factors that have been shown to influence clinical studies on pain and analgesia are sex and age differences in the likelihood of self-reporting pain in a clinical setting, and the subjective nature of pain (Ferrell et al., 1991; Reddy et al., 2012; Dampier et al., 2013).

### Sex and age differences in vlPAG MOR expression and binding

The PAG is a critical neural site for opioid modulation of pain (Reynolds, 1969; Behbehani and Fields, 1979; Morgan et al., 1991; Morgan et al., 1992; Zhang et al., 1998; Loyd et al., 2008). In the present study, using both immunohistochemistry and autoradiography, we report that aged rats exhibit reduced vlPAG MOR expression and binding compared to adults. This suggests that the age-induced decrease in morphine potency is driven, in part, by diminished vlPAG MOR. Previous studies have also reported an attenuation in MOR in the frontal cortex and striatum of aged male rats compared to adults (Messing et al. 1981; Hess et al., 1989). Reduced MOR binding in the midbrain and thalamus of aged female rats has also been shown (Messing et al., 1980). Together, these studies suggest that the reduction in MOR observed in aged rats is not specific to the PAG. Importantly, we did not find a significant reduction in MOR in the inferior colliculus as a function of age or sex, suggesting that these age-associated changes may be limited to CNS sites implicated in pain and reward. No reduction in neuronal number within the vlPAG was observed, indicating that the observed reduction in MOR was not due to overall neuronal loss.

No impact of persistent inflammatory pain on MOR expression or binding in the vlPAG was noted. Similarly, Thompson et al. (2018) reported no change in MOR availability or expression in the PAG of male rats following induction of neuropathic pain, although decreased MOR was observed in the caudate putamen and insula (Thompson et al. 2018). In contrast, Zollner et al. (2003) reported increased MOR binding in the dorsal root ganglia following intraplantar CFA (Zollner et al., 2003), suggesting that the effects of persistent inflammatory pain on MOR are region-specific. Clinical studies utilizing PET scans to assess for changes in MOR binding reported that individuals experiencing rheumatoid arthritis pain exhibit decreased MOR binding in the straight gyrus and the frontal, temporal, and cingulate cortices (Jones et al., 1994); in contrast, individuals experiencing fibromyalgia pain exhibited reduced MOR binding in the nucleus accumbens, amygdala, and cingulate cortex (Harris et al., 2007). Neither of these studies reported an impact of pain on MOR binding in the PAG. A third study examining MOR binding in individuals experiencing post-stroke neuropathic pain (centrally-versus peripherally-localized) reported reduced PAG MOR binding in individuals experiencing neuropathic pain localized centrally not peripherally, compared to controls (Maarrawi et al., 2007). Together, these results suggest that PAG MOR expression and binding may be altered as a function of the pain location or modality.

In the present study, we found no impact of sex on vlPAG MOR expression. These results are contradictory to our previous findings in which we reported significantly higher levels of vlPAG MOR in adult males than females (Loyd et al., 2008). In that study, sex differences in MOR were primarily driven by diestrus females, with smaller, non-significant differences noted for the other stages of estrous in comparison to males. In the present study, although the estrous stage was determined, it was not included as a factor of analysis due to a lack of power. Methodological differences between the two studies may also have contributed to these conflicting results, including differences in antibody specificity and 3-dimensional confocal versus 2-dimensional light-field microscopy image acquisition and assessment of MOR density.

The finding of no significant effect of sex on vlPAG MOR expression or binding suggests that the observed sex differences in morphine potency are not driven by vlPAG MOR expression or binding as previously proposed (Loyd et al., 2008). Recent attention has focused on sex differences in neuroimmune signaling, and in particular, microglia, as a contributing factor to morphine’s dimorphic effects (Sorge et al., 2011; Rosen et al., 2017; Doyle and Murphy, 2017; Eidson et al., 2019; Eidson and Murphy, 2019). In addition to binding to neuronal MOR, morphine has been shown to bind to the innate immune receptor toll-like receptor 4 (TLR4), localized primarily on microglia (Hutchinson et al., 2007; Hutchinson et al., 2008; Hutchinson et al., 2010). Morphine action at TLR4 has been shown to decrease both glutamate transporter (GLT1 and GLAST) and GABA_A_ receptor expression, and upregulate AMPA receptor, resulting in an overall increase in neural excitability (Song and Zhao, 2001; Holdridge et al., 2007; Wang et al., 2012; Eidson et al., 2017; Ogoshi et al., 2005; Stellwagen et al., 2005). As morphine works primarily via hyperpolarization (Chieng and Christie, 1996), this increased neural excitability within the vlPAG directly opposes morphine action (Ingram et al., 1998). Our lab has recently reported that blockade of vlPAG TLR4 signaling via (+)-naloxone results in a 2-fold leftward shift in the morphine dose response curve in females, but not males (Doyle et al., 2017). Further, we report that pathogenic activation of TLR4 via systemic administration of the bacterium lipopolysaccharide (LPS) results in a significantly higher vlPAG expression of the pro-inflammatory cytokine IL-1ß. Reduced vlPAG expression of the anti-inflammatory cytokine IL-10 was noted in females, but not males, following LPS. Increased levels of neuroinflammation have been reported in aged animals suggesting that changes in neuroimmune signaling may also contribute to the reduction in morphine efficacy (Gorelick et al., 2010; VanGuilder et al., 2011; Norden and Godbout, 2013).

### Summary

Despite the recognition that undermanaged pain in the elderly is a growing clinical problem, remarkably little basic research has been conducted. The results of the present study establish that morphine’s ability to attenuate persistent inflammatory pain is reduced in aged male rats, likely due to reduced density and functionality of vlPAG MOR. Together, these results suggest that PAG MOR may provide the anatomical basis for observed age differences in morphine analgesia, and present PAG MOR (OPRM1) as a potential target for gene therapy in the elderly.

## Author Contributions

EF and MR contributed equally to this work. AZM, MR and EFF designed the research and wrote the paper. EFF, RIH, MK, and MR performed the research and analyzed the data.

---

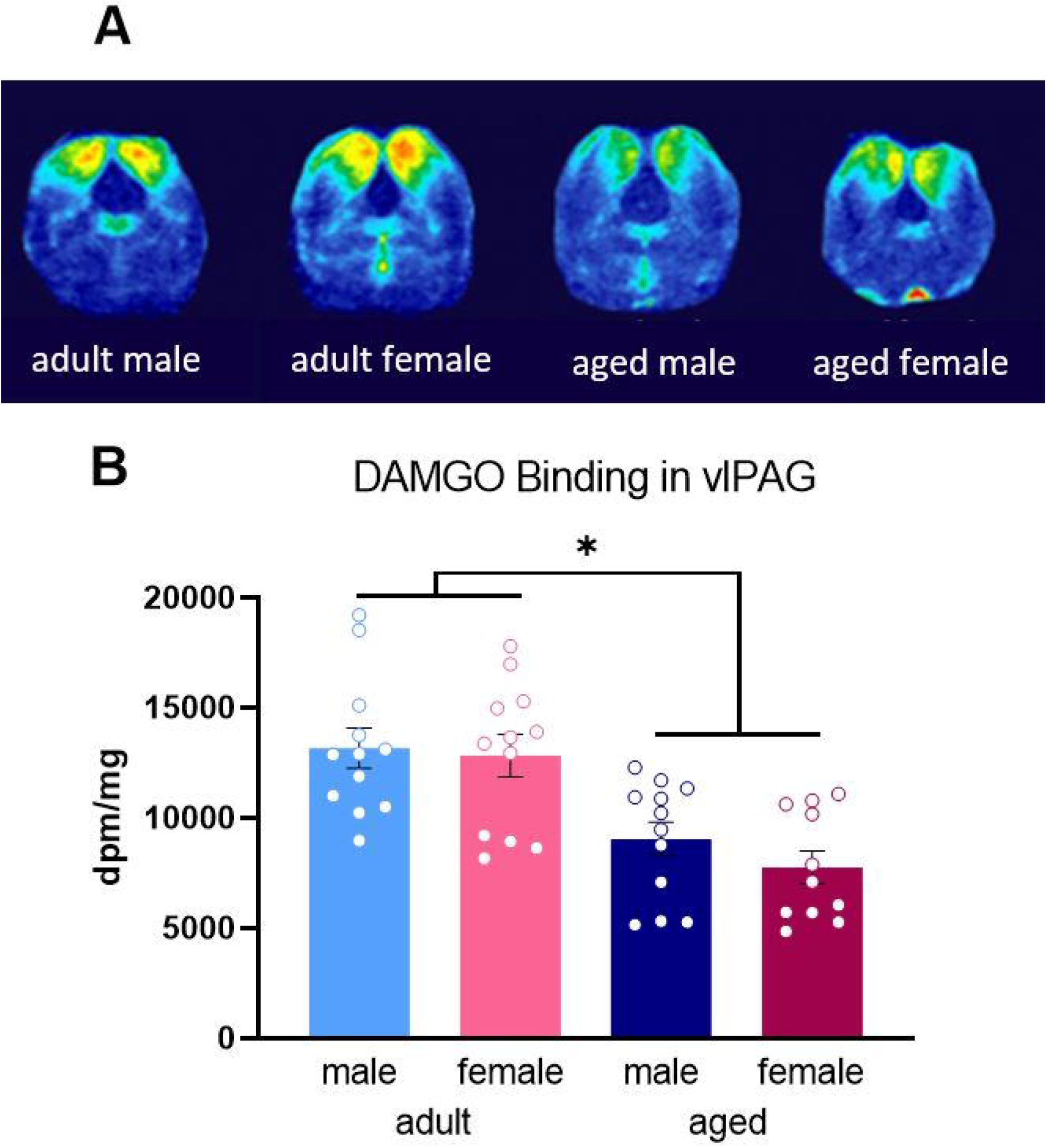

